# Collective intelligence facilitates emergent resource partitioning through frequency dependent learning

**DOI:** 10.1101/2024.03.01.582983

**Authors:** Mina Ogino, Damien R. Farine

## Abstract

Deciding where to forage must not only account for variation in habitat quality, but also where others might forage. Recent studies have suggested that when individuals remember recent foraging outcomes, negative frequency-dependent learning can allow them to avoid resources exploited by others (indirect competition). This process can drive the emergence of consistent differences in resource use (resource partitioning) at the population level. However, indirect cues of competition can be difficult for individuals to sense. Here, we propose that information pooling through collective decision-making—i.e. collective intelligence—can allow populations of group-living animals to more effectively partition resources relative to populations of solitary animals. We test this hypothesis by simulating (i) individuals preferring to forage where they were recently successful, and (ii) cohesive groups that choose one resource using a majority rule. While solitary animals can partially avoid indirect competition through negative frequency-dependent learning, resource partitioning is more likely to emerge in populations of group-living animals. Populations of larger groups also better partition resources than populations of smaller groups, especially in environments with more choices. Our results give insight into the value of long- vs. short-term memory, home range sizes, and the evolution of specialisation, optimal group sizes, and territoriality.

## Introduction

Animals share the environment with other animals of the same and different species, and the resulting direct and indirect interactions can shape community structure and composition [1–3]. A key question is what factors facilitate the coexistence of animals that compete over the same set of resources [4–7]. Active exclusion of competitors from the area or preventing competitors from consuming the resource through direct interaction (contest competition, e.g. establishing territories or agonistic interactions [8, 9]) is one way of securing required resources for survival and reproduction [10, 11]. However, such direct interaction can be energetically costly, risky and/or inefficient [10]. As a result, animals often instead compete indirectly, by exploiting resources before others (i.e. scramble, exploitative or indirect competition) and avoiding resources that are already heavily exploited (e.g. resource use specialisation [12–14] or the ideal free distribution [15]). Recent studies have demonstrated the potential for indirect competition to shape ecological communities (e.g. by driving selection for mutualisms [16, 17]) and individual niche specialization (e.g. [18, 19]), but much less is known about its role in shaping within-species social structure ([19]; but see [20]).

Specialisation plays a fundamental role in shaping ecological processes and various behaviours at different (time and ecological) scales ([21–23]; e.g. [24, 25]). Classical ecological theory posits that reduced resource overlap (stronger resource partitioning) between species allows increased (net and individual) resource use and the coexistence of diverse species in the same environment [26–28]. Such resource partitioning is often thought to emerge through frequency-dependent interactions [29–31], with the resulting specialisation in resource use eventually (over generations) leading to shifts in morphological traits between species [18, 32–34], coevolution of morphological traits and food resources [35], or adaptive radiation [36]. These enable species to efficiently access a set of resources, leading to niche specialisation. For example, populations of two salamander species, *Plethodon cinereus* and *P. hoffmani*, have greater segregations in prey types, and their trophic morphological traits significantly diverge, in sympatry relative to allopatry [37]. Within generation time, species can also shift their resource use and partition resources depending on the composition of species sharing the same environment. For example, a population experiment in *Anolis* lizards showed that stronger resource partitioning between species corresponded to lower competition among these species [38]. When sharing the environment with a species (*A. gingivinus*) whose resource use was highly overlapping (i.e. between-species competition was strong), *A. wattsi* caused changes in resource uses (prey size and perch height to use) and lowered growth rates of the sympatric *Anolis* species. However, when sharing the environment with a species (*A. bimaculatus*) whose resource use was less overlapping, the presence of *A. wattsi* did not have fitness consequences on the sympatric *Anolis* species. Such patterns of differentiation in response to competition can lead to greater net exploitation of resources. For example, in pollinator species, the more species are present in the environment, the more resources the community uses (and more seeds were produced) because each pollinator species specialises in a limited range of resources and resource use overlaps less between species, meaning that a broader range of plants are used in total [24]. Thus, specialisation in resources through frequency-dependent interactions allows species to coexist in the same environment, by reducing competition among species [21, 38, 39].

Resource partitioning can also exist among conspecifics, with intraspecific variation in resource use being linked to differences in morphology or behaviour between individuals (e.g. according to sex through ecological sexual dimorphism [40], developmental stage following ontogenetic niche shift [41], or personality [42]). Energy requirements—or access to a limited pool of resources—can drive sexual dimorphism and corresponding resource partitioning between sexes [25, 35, 40, 43]. For example, in sooty oystercatcher *Haematopus fuliginosus* [40] and hummingbirds *Eulampis jugularis* [25, 35], sexual dimorphism in bill morphology has been linked to males and females specialising in different food types, thereby reducing competition over limited resources within breeding pairs, enabling them to maintain close spatial proximity or maintain smaller territories. Such intraspecific variation in specialisation can initially emerge and be maintained through frequency-dependent selection that persists across generations. For example, experimental populations of initially cadmium intolerant fruit flies *Drosophila melanogaster* evolved and expanded their niche to cadmium- laced resources which fewer individuals competed over [42]. Thus, in both intraspecific contexts and in broader ecological communities, we can observe extensive resource partitioning—whereby individuals or species have evolved to specialise in distinct types of resources—through frequency-dependent mechanisms.

Frequency-dependent learning can also play a major role in shaping how animals distribute themselves over resources at much finer timescales. By sampling the environment, individuals can detect resource gradients that arise from exploitation by others [44], thereby allowing them to choose resources corresponding to their availability (the ideal free distribution hypothesis [45]). If resources are temporally stable, the repeated use of information about what resources are under-utilised (i.e. negative frequency-dependent learning) can lead to the emergence of specialisation in resource choice. An experiment in house sparrows (*Passer domesticus*) demonstrated this process by introducing naïve individuals into captive groups of specialists where individuals had an already established specialisation in sets of foraging cues [32]. Through a mix of social and individual learning, newly introduced naïve individuals learned the cues that specialists used, as well as the alternative cue that had the same starting food availability but was less exploited by specialists. Over time, previously naïve individuals established a preference for the alternative cues because it generated a higher payoff (as the initial specialists foraged less on them, i.e. negative frequency-dependent learning). Such specialisation, driven by individual-level decisions, can potentially scale up to population-level outcomes. For example, GPS tracking of northern gannets (*Morus bassanus*) from multiple colonies showed that neighbouring colonies maintained different foraging areas, driven by individual-level density-dependent competition both within and between colonies (the density-dependent hinterland model [34], an extension of the ideal free distribution hypothesis [45]). Thus, by remembering where they were recently successful, individual animals can make foraging decisions that ultimately avoid indirect competition and maximize their ability to extract resources from the environment. In this way, negative frequency-dependent learning can lead to establishing specialisations in where they foraging (space use) [46], whereby individuals maintain distinct preferences for particular foraging locations that results in repeatable within-individual space use and consistent differences in space use among individuals.

Previous studies investigating the role of frequency-dependent learning and indirect competition in shaping the foraging strategies—and the emergence of resource partitioning— have generally focussed exclusively on individual-level decisions, and how these scale up to behavioural specializations within a social group [25, 47] or between populations [34]. While in many group-living animals individuals make their own decisions about where to go or what to do (e.g. some seabirds live in colony, but foraging trips are made by individuals or small groups), other animals live in stable and cohesive groups (e.g. some groups of primates, other mammals, and birds; see [48, 49]), meaning that decisions are made by aggregating up individual-level preferences to produce one group outcome. While conflict resolution in preferences among group members represents a challenge to decision-making [26, 48, 50], having more than one individual contributing to decisions also allows groups to pool different information that individuals have sensed from the environment, giving groups the opportunity to benefit from emergent collective intelligence [51]. Information pooling is a process whereby the noise that is inherent in making estimations is averaged out when pooled (or averaged) across individuals. For example, in 1907 Galton asked 787 visitors at an agricultural fair to guess the weight of an ox, finding that the mean of participants guesses was within 1% difference from the true weight, despite the individual guesses themselves being much less accurate [52]. Collective intelligence can then arise when the interactions among group members allows the group to express properties that individuals do not [53, 54]. Combining information pooling with collective intelligence—for example when making collective decisions about where to go—can allow groups to be much better at detecting information from noisy signals than solitary individuals can.

Widespread evidence, from a range of taxa [54, 55], suggests that individuals can benefit from collective intelligence when groups aggregate individual preferences and make decisions by selecting the preference representing the majority of group members. For example, in colonies of honey bees *Apis mellifera* and ants *Leptothorax albipennis*, individuals collect information and recruit group members to potential nest sites [55]. Through repeated recruitment interactions, group members pool information about the quality of potential locations and collectively make decisions for the new nest sites following a quorum or majority rules. Crucially, the collective decision-making process allows groups to make accurate decisions effectively without all of group members having to sample all candidate sites or for individuals needing to carefully assess the potential nest sites. Thus, collective intelligence is particularly powerful when the ability to extract information from the environment is limited or costly to obtain. This ability might be particularly important when making experience-based decisions as a way of avoiding resources used by conspecifics (i.e. negative frequency-dependent learning). While long-term memory may be useful for making decisions in environments that change relatively slowly (e.g. with seasonality), the social environment can produce much faster resource dynamics, thereby increasing the potential value of recent experiences. However, these are, by definition, rarer. Thus, collective behaviour may allow animal groups to more effectively aggregate fewer, but more recent, experiences into decisions that result in better avoidance of others.

In this paper, we integrate the concepts of information pooling and collective decision-making with resource partitioning through negative frequency-dependent learning. We propose a process (herein, ‘collective sensing of indirect competition’) where collective intelligence can allow cohesive groups to effectively estimate (or ‘sense’, using negative frequency-dependent learning) resource gradients created by the foraging decisions of others without requiring any direct intergroup interactions. We hypothesise that groups should be better than solitary individuals at collectively sensing modifications of resources by competitors. Sensing cues about where others have foraged can then underpin the emergence of partitioning of, or specialisation in, resource use at the population level. Our hypothesis predicts this should result in a greater exploitation of resources by populations of group-living animals than what is possible in solitary decision-makers. Specifically, we predict (i) groups develop stronger patch preferences than solitary individuals do and (ii) individuals in groups successfully forage more frequently than solitary individuals.

To test our hypothesis, we simulate environments consisting of populations of cohesive groups of the same size (group size varies across simulation from 1 to 93, herein the term ‘groups’ includes solitary individuals) that repeatedly and simultaneously choose which patch to forage at. Our simulations are designed such that all individuals in a simulated population get an equal payoff if groups all choose different patches from one another. If multiple groups select the same patch, the resources at a patch are split equally among these groups and randomly assigned to group members (without any cues or cost of contest competition, between and within groups). At each timestep, individuals first decide on what patch they prefer to forage, based on where they were recently successful (using short-term memory), and groups then collectively choose one patch using a majority rule (meaning that some group members may not get to forage at their chosen patch). Further, to explore if and how spatial structure (e.g. habitat structure or restricted home ranges) can facilitate the emergent differences in patch preferences, we limit the access of each group to a specific subset of contiguous patches. We also explore different ecological scenarios (e.g. resource rich vs. poor environments) and the relative value of shorter-term vs. longer-term memory. In summary, our model examines if groups can avoid competitors (other groups that share preference for the same patch) using only the information based on recent foraging success (negative frequency-dependent learning), and whether collective intelligence allows populations of groups to be more effective at doing so than populations of solitary individuals. Our study therefore provides new insights on the role of collective intelligence in the evolution of resource partitioning, and we discuss the implications of our model for understanding optimal group sizes and the evolution of territoriality.

## Methods

### General model structure and simulations

We simulate individuals having to make repeated decisions about where to forage while living in an environment in which resources can be exploited by other individuals (Figure 1). Populations consist of either solitary individuals (herein solitary individuals have a group size, N, of 1) or groups (we always use odd-sized groups to increase the chances of having clear majorities), and all groups in the population have the same size. Each group can access all patches in the environment, the number of patches (N_P_) is equal to the number of groups (N_G_), and the number of resources within each patch is equal to the group size. Patches are fully replenished in each timestep. Thus, the total amount of resources matches the number of individuals in the population (but we also explore the consequences of resource-poor environments in the Supplemental Information sections 1 & 2).

**Figure 1.**
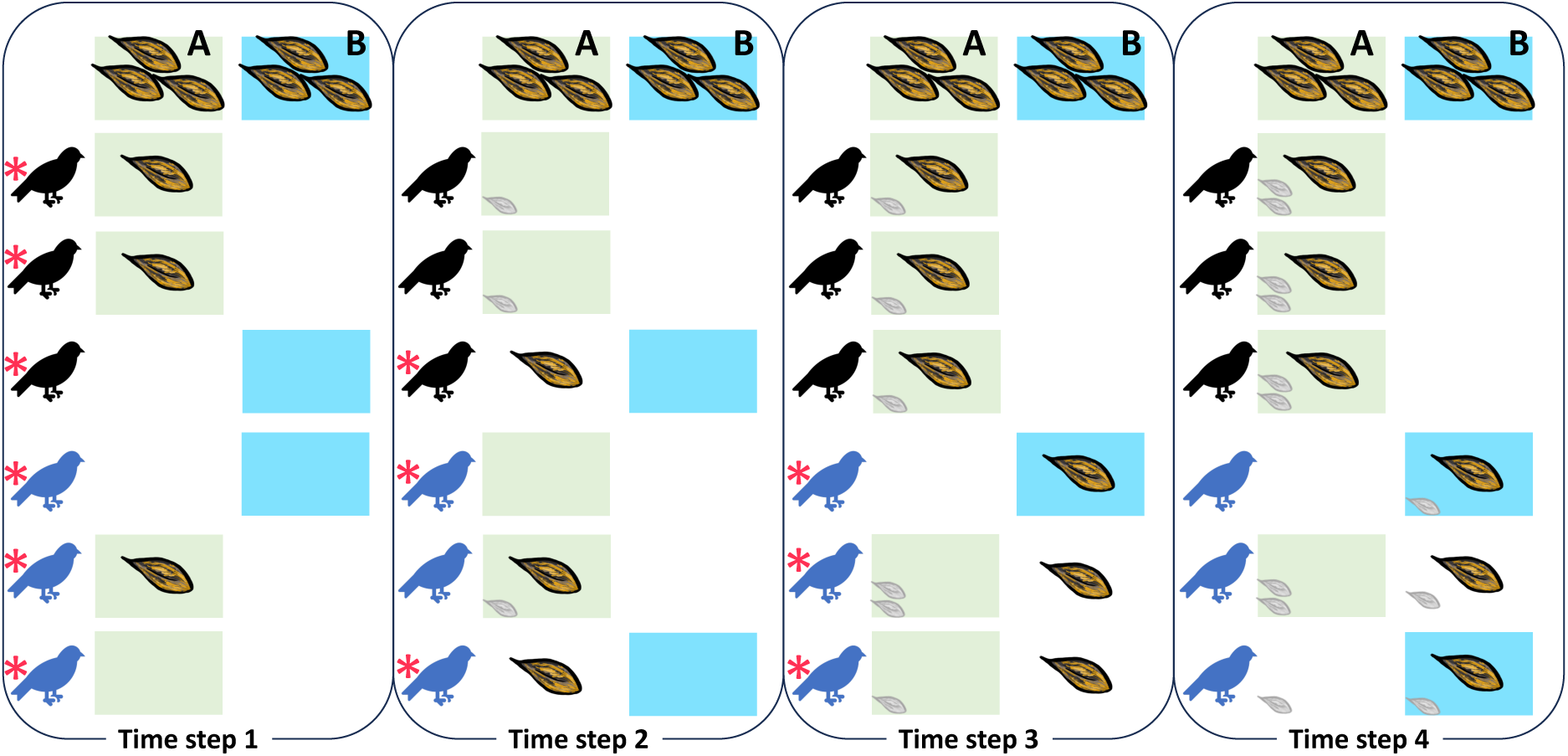
Example showing the implementation of individual preferences, collective decisions, and allocations of rewards at each timestep in the simulation. Here, two groups (black and blue), each consisting of three group members, are shown. In timestep 1, all individuals randomly (indicated by *) select between patch A (choices for patch A are marked by a green background) and B (choices for patch B are marked by a blue background), as individuals have no prior information. By chance, both groups select patch A (the majority of group members had green background at timestep 2), so the three resources available at that patch (shown as big seeds) are split into two (group 1) and one (group 2), and then randomly allocated to individuals in each respective group. At the beginning of timestep 2, three individuals (across the two groups) that were rewarded at timestep 1 (the total reward from three previous steps for each individual is shown as shaded small seeds) each show a preference for patch A, resulting in a majority for group 1 selecting patch A. In this case, group 2 also again selects patch A, and thus the rewards are again shared between groups. In the third timestep, groups both establish majority of preferences for patch A, but group 2 makes an explorative choice (*; occurring by default in 1% of decisions; *exploration*=0.01). As all individuals are rewarded in timestep 3, individuals in group 2 start developing preferences for patch B. By being rewarded in patch B at timestep 4, the members of group 2 then continue to select patch B as their preferences, whereas the members of group 1 continue selecting patch A. This is just one scenario by which differences in patch preferences can emerge among groups (and not all scenarios require explorative decisions).

In each timestep, individuals first select a patch based on their recent history of foraging success (the choice is initially random as individuals have no history, and is random if they have not been successful in the history captured by their memory). When individuals are in groups with N > 1, they then use a majority rule, foraging at the patch that the largest number of group members preferred (selecting at random between most-preferred patches if there was not one clearly preferred patch). When foraging at the selected patch, each individual receives one resource if the group is the only one to have selected that patch. If more than one group selects the same patch, resources are equally split (or as close as possible) among groups and randomly allocated among the group members. This simulates indirect competition as we do not include any costs of contest competition (e.g. agonistic interactions), it instead assumes that individuals from two groups selecting same patch would be rewarded on average half the time, and removes any order of arrivals effects. While we could have selected one random group to have all group members rewarded, we decided *a priori* that this would be too likely to generate strong differences in rewards, and therefore future preferences (for that patch versus other patches), among groups. By contrast, our approach means that only a subset of individuals within each group gain information that other groups have selected the same patch, making the simulation more realistic. Finally, we note that an individual can only access information about its own previous foraging history (prior successful foraging events at each patch) within its memory capacity, and this foraging history is only used for future patch choices by the individual. Individuals have no information about the identity of other groups, frequency that they have foraged at different patches, or rewards that other group members (of their group or of other groups) that have led to establishing their preferences in a given simulation round.

Individuals’ choices of patches in each timestep are determined in two steps. First, individuals estimate the quality of each patch based on their prior experience (number of rewards) within a given capacity of memory (we set this as *history=3*, taking account three previous timesteps, but also explore the effects of the *history* parameter in the Supplemental Information Section 3). Patch quality is estimated using 1/{1+*e^-rewards+sensitivity^*}, where *sensitivity*—a measure of how much previous experience affects the estimate of quality—is set as 3. Larger values of *sensitivity* correspond to a stronger contribution of successful patch visits to estimations of quality (and, correspondingly, a smaller estimation of quality for patches where *rewards*=0). Finally, individuals select a patch based on these quality estimates by setting the probability of selecting a patch to be linearly proportional to the quality. Second, when in a group, groups then use a majority rule when deciding where to forage. Note that preferences are only temporary, for the purpose of selecting one patch based on previous rewards, and are not observed by others or remembered in future decisions. Groups also occasionally select an accessible patch at random (*exploration*; default is 1% of decisions) to mimic exploration of the environment (the effects of the different exploration rates are reported in the Supplemental Information section 4).

Simulations ran for 300 timesteps, and we repeated simulations 100 times for each combination of parameter values and environment types (see below). Additionally, we test if simulations remain stable if the population starts from non-random state, here where groups start in a state where they partition resources at the highest rate possible from the simulation outcomes (see Supplemental Information section 6).

### Incorporating spatial restrictions into the model

In our model (*herein* full network), all groups can access all patches (Figure 2*a*). However, in reality animals are limited in where they can go by physical aspects of their environment [56]. As a result, individuals or groups that are closer to one-another are more likely to interact [57] through shared space use. To capture this process, we also include models where each group is limited to a subset of patches, and where subsets overlap with other, neighbouring groups. Specifically, all groups can access three neighbouring patches, which we call a ring network (Figure 2*b*, note that this is technically a bipartite network), capturing the limited space use of animal groups. We investigate the effects of increasing the number of patches that groups can access in Supplemental Information section 5.

**Figure 2.**
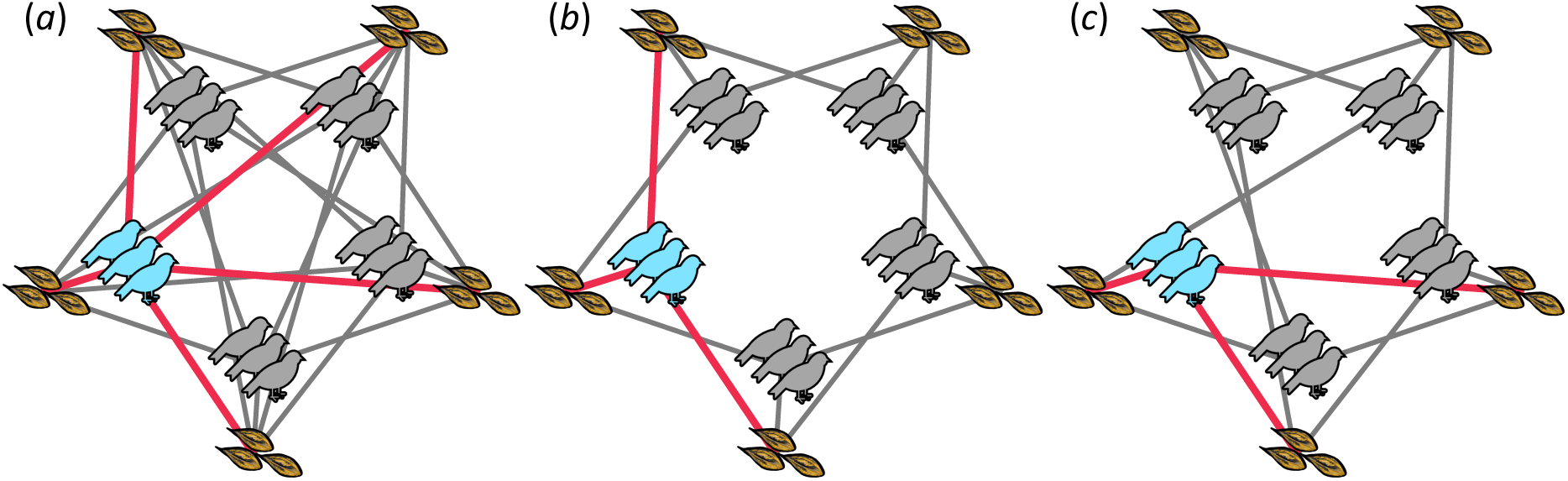
Visual representations of the patches accessible to each group in different environments. Here, five groups, each consisting of three individuals, live in an environment containing five patches (the number of patches N_P_ is always equal to the number of groups N_G_) each containing three resources (all patches have resources equal to the group size). In the first environment type (full network, *a*) all groups can access all of the possible patches (N_P_=N_G_= 5, lines represent which patches each group can access). The thicker red lines show the patches accessible by one group (here the light blue group), with the same pattern repeated across all groups. In the second environment type (ring network, *b*) groups are spatially restricted (e.g. due to a limit in their ability to range) to access a limited subset of the patches (default N’_P_=3; where N’_P_ is the number of patches that the group can select from). Accessible patches are contiguous (e.g. fall within a restricted home range), and thereby neighbouring groups overlap in being able to access a similar subset of patches. Finally, to examine the effect of limited accessibility (i.e. the number of patches available) versus spatial structure (e.g. home ranges), we also implement a random network model (*c*) in which groups can access a fixed number of patches (default N’_P_=3), but these are randomly distributed in relation to other groups.

Because both changes in the extent of connectivity between patches and groups (i.e. the number of patches groups can access) and the pattern of connectivity among groups (the structure of the network of potential indirect competition) could impact the development of patch preferences, we also implement a random network (Figure 2*c*). To create the random network, we start with each ring network and first randomly select one edge connecting one group to a resource patch. We then swap the edge to a new patch, such that the targeted patch is not one already available to that group (i.e. overlapping with an existing edge). We repeat this process a number of times, equivalent to half of the number of edges present in the network, to generate more random connections between groups and patches. As a result, in both the ring and random networks, groups can access a fixed number of patches, but while groups overlap in their available patches only with neighbouring groups in the ring network, they potentially overlap with a larger (or smaller) number of groups in the random network (Figure 2).

### Quantifying patch preferences, resource partitioning, payoffs, and conflicts within the group

We evaluate if, and how animals can detect and avoid indirect competition, and the fitness consequences arising from this, by recording four sets of metrics: (i) the tendency for groups to express a preference for a given foraging patch over time and (ii) the distribution of groups across patches (resource partitioning) within each timestep. We also (iii) quantified the number of resources gained by individuals (as a measure of how well distributed groups are over the available patches) and (iv) the extent of conflict (or not) in preferences within each group as measures of fitness (i.e. a cost that arises from living in a group [48]). We record each of these metrics over the final 30 timesteps (the final 10% of the simulation timesteps) of each simulation. We define the expressed patch preference and resource partitioning using the normalised entropy. Entropy (Shannon entropy [58]) is calculated using 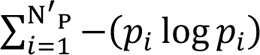. For patch preferences (a within-group measure), we set *p_i_* as the proportion of the last 30 timesteps that the group used each patch *i*. For resource partitioning (a between-group measure), we set *p_i_* as the proportion groups in each timesteps that are using each patch *i*. We divide the entropy measure by log N’_P_ to normalize the metric, because the possible maximum values for the entropy measure changes corresponding to the number of patches available to groups (or the number of patches in the environment, N_P_, in the case of resource partitioning). To convert entropy to have higher values for greater patch preferences, we subtract entropy from 1. Thus, patch preferences can range from 0 (groups use a different patch each timestep) to 1 (groups only use one patch in all timesteps), while resource partitioning can range from 0 (only one patch is used by all groups in a given timestep, i.e. no resource partitioning) to 1 (every patch is used by one group in a given timesteps).

We define individual foraging success (i.e. foraging rates) at individual level as the average proportion of timesteps that individuals acquired a resource. This could in theory range from 1/N_G_ (where N_G_ is the number of groups in the population) if all groups always select the same patch to 1 if all groups perfectly distribute themselves over the available patches. The baseline expectation is higher when groups have access to a limited number of patches (i.e. N’_P_ < N_G_).

Finally, we capture information about consensus and conflict in patch choices within groups (within each timestep) as the agreement among group members. We define agreement as the percentage of group members that express the same choices as the patch most-commonly selected among group members. This measure could range from 1/N’_P_ if individuals are always perfectly distributed in their patch preferences across each timestep to 1 if all individuals always agree on selecting the same patch in all timesteps.

### Parameterising the simulations

We ran simulations spanning a range of different environment types (full, ring and random networks), group sizes (discontinuously from 1 to 93 individuals per group; a total of 37 different group sizes per condition), and number of groups and patches (7, 15, 31 groups or patches). In the main text, we present the results from setting the number of accessible patches that groups can forage (for ring and random networks) at N’_P_=3, *history*=3, and *exploration*=0.01, and always keep the number of resources equal to the number of individuals in the population (N_P_=N_G_). As environments can sometimes be resource poor, we also explored the effect of reducing the total amount of resources by reducing the total number of patches (Supplementary Information Section 1) and the number of resources within each patch (Supplementary Information Section 2). We also explore the effect of increasing memory (*history*=7) and exploration rate (*exploration*=0.1 and *exploration*=0.25), and increasing the number of patches that groups can forage at in the ring and random networks (N’_P_=5).

To quantify the effect of each parameter (total amount of resources and resource distributions, *history*, *exploration*, N’_P_) on patch preferences and resource partitioning, we calculate ‘the tendency to express higher metric value’. We do this calculating the proportion of simulation outcomes where patch preferences are equal to or above 0.5 and resource partitioning is equal to or above 0.9 (these values correspond to transitions between low and high values, and closely match the median values across simulation parameters). This ‘tendency’ captures the probability that the outcome of the simulations is in the upper half of the distribution of the outcomes across parameter sets, and enables us to visualise the effect from each parameter on patch preferences and resource partitioning.

### Estimating the null expectation under random decision-making

We expect the baseline values of our metrics to differ depending on simulated conditions (i.e. parameter values, e.g. the number of groups N_G_ or accessible patches N’_P_). For example, when groups can access fewer patches (in the ring or random networks), then they should have higher values in terms of preferences and average rewards even if they select patches randomly. Thus, for each parameter set we create a corresponding reference model in which we simulate random, or uninformed, collective decisions (*history*=0, which also corresponds to *exploration*=1) while keeping all other parameters the same as the simulation, thereby estimating the baseline (or null) expectation for each simulated condition.

## Results

### Larger groups develop stronger patch preferences

Both solitary individuals and groups can show consistent preferences for one patch when making decisions based on individuals’ previous experience (short-term memory), relative to selecting patches randomly (i.e. *history*=0; Figure 3). These results are consistent and not sensitive to parameter choices (Figures S3.1a, S4.1a, S4.3a, S5.1a). Larger groups also develop stronger preferences than smaller groups or individuals. The shape of this relationship is non-linear, with an interaction between the environment (the number of patches accessible to each group, N_P_) and group size. Specifically, smaller- and intermediate-sized groups in full networks develop weaker preferences compared to groups of the same size in ring and random networks, with solitary individuals showing almost no preferences (i.e. only marginally more consistent use of a single patch than random patch uses, i.e. *history*=0, Figure 3c). As the number of patches available to each group increases, the ability for groups to develop a preference weakens (see also Figure S5.1a). Conversely, when groups are restricted in the number of patches available (ring and random networks), they develop stronger preferences, even at small group sizes. Groups develop stronger patch preferences, when the memory capacity is larger (Figure 4a), when they make fewer exploratory decisions (Figure 4b), and when there are fewer patches to choose from (Figure 4c).

**Figure 3.**
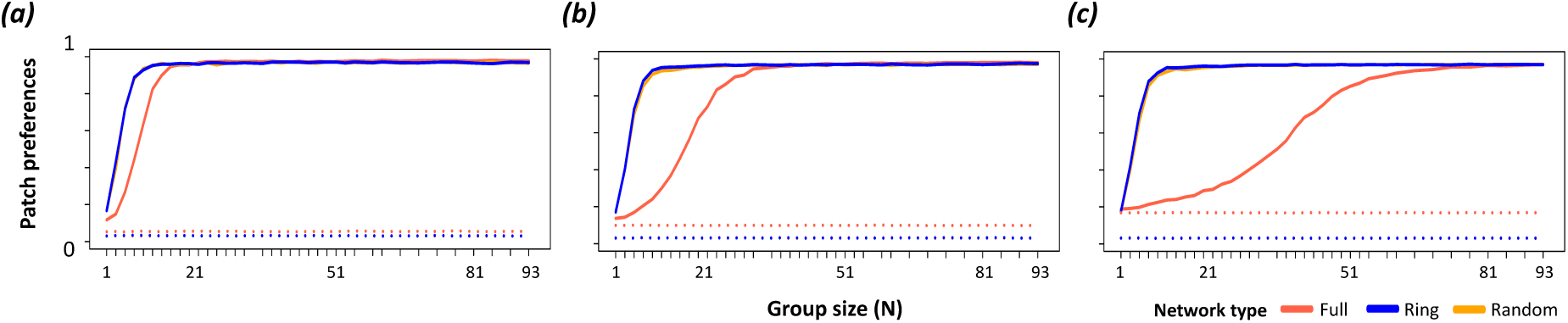
Groups can develop strong patch preferences. These plots show patch preferences (1-normalised entropy in patch use within groups) over the last 30 timesteps in simulations with (a) 7, (b) 15, and (c) 31 groups per environment (N_G_; N_G_=N_P_). Patch preference is higher in populations of groups than populations of solitary individuals, and increases with group size (x axis) when individuals use prior experience (*history*=3, solid lines) to choose foraging patches (relative to making uninformed patch choices at all timesteps, dotted lines, *history*=0). While only large groups develop preferences when they can access many patches [e.g. N_P_=31 in the full network, orange line, in (c)], even small groups (e.g. N=11) can develop strong preferences when they are restricted in the number of patches that they can access (i.e. ring and random networks, where N’_P_<N_G_). Note that the maximum patch preference is generally slightly less than 1 because groups make explorative decisions in 1% of timesteps (*exploration*=0.01). Colours show full (N’_P_=N_P_), ring (N’_P_=3), and random (N’_P_=3) networks (note that ring and random networks largely overlap and may not both be visible).

**Figure 4.**
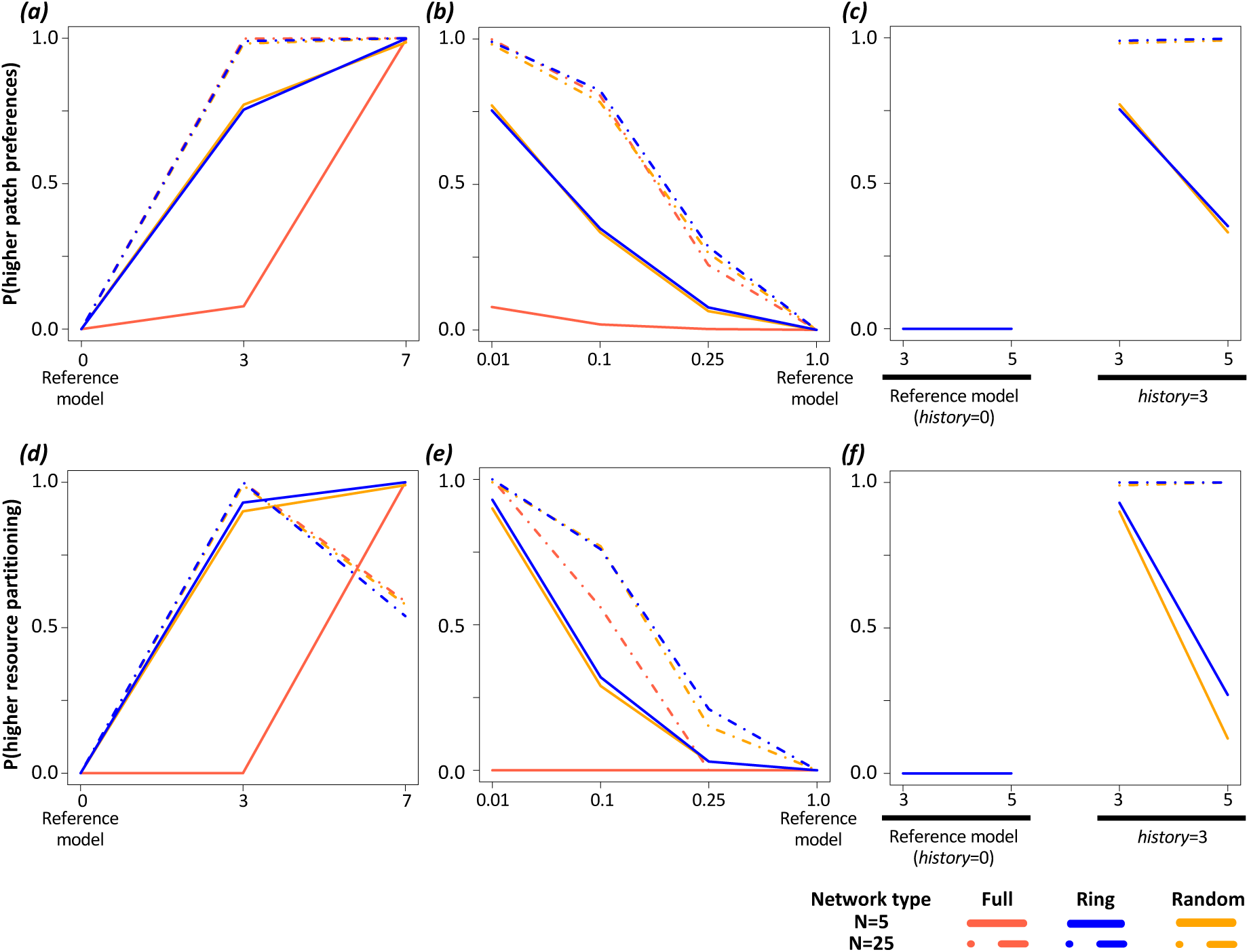
Memory, rates of explorative decisions, and the number of accessible patches all affect the establishment of preferences and the emergence of resource partitioning. Plots show the effects of memory capacity (*history*; a,d), the frequency of explorative decision-making (b,e), and the number of accessible patches (c,f) on patch preferences (top row) and resource partitioning among groups (bottom row). The y axis shows the tendency of the simulation with the given parameter set to express a higher metric value that is higher than or equal to the median of the default parameter set (see methods). Colours show full (N’_P_=N_P_), ring, and random networks (note that results for the different networks often overlap with each other). All plots except ones showing the effects of the number of accessible patches (c,f) include all network types. Full results for different history, exploration rates, and number of accessible patches are shown in Supplemental Materials sections 3, 4, and 5, respectively.

### Populations of intermediate-sized groups better partition resources

On average, populations consisting of groups are better at partitioning resources than populations consisting of solitary individuals, and as a result groups are more often successful at acquiring resource than solitary individuals (Figure 5, Figure S3.2, Figure S4.2, Figure S4.4, Figure S5.2). Importantly, while individuals in larger groups can substantially outperform making random decisions (*history*=0), solitary individuals (group size: 1) can only slightly outperform the baseline (null) expectation.

**Figure 5.**
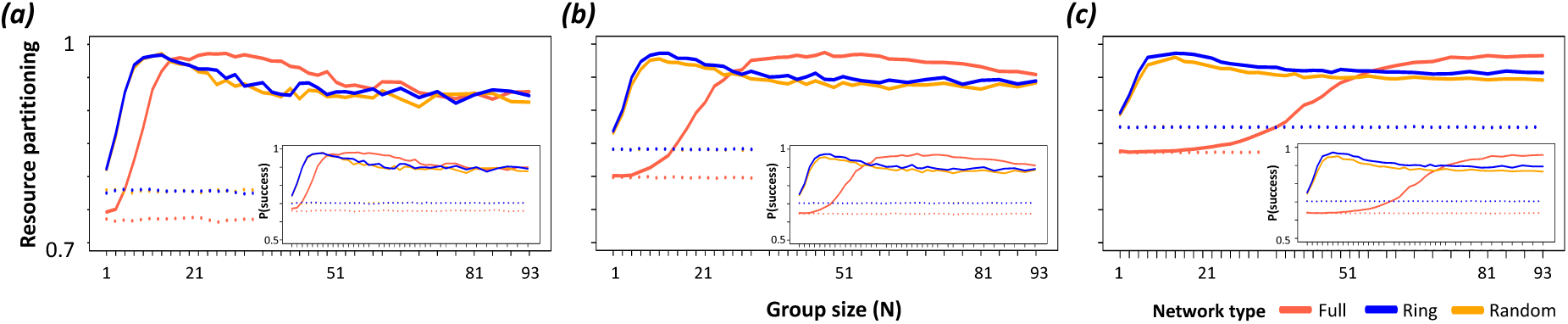
Populations of groups show greater resource partitioning, with individuals in groups experiencing higher foraging rates. Plots show the resource partitioning (normalised entropy in patch use across groups) and the probability that an individual is successful in acquiring a resource (insets) over the last 30 timesteps in simulations with (a) 7, (b) 15, and (c) 31 groups per environment, N_G_. Populations of larger groups express greater partitioning of resources among groups than populations consisting of solitary individuals, with the latter only slightly outperforming populations where individuals make uninformed choices (dashed lines). Greater partitioning corresponds closely with higher foraging rates experienced by individuals. Colours show full (N’_P_=N_P_), ring (N’_P_=3), and random (N’_P_=3) networks (note that ring and random networks largely overlap).

While larger groups can develop stronger patch preferences (Figure 3), resource partitioning and foraging success peaks at intermediate group sizes (Figure 5). The location of this peak shifts in resource poor environments, where patch sizes are smaller than group sizes (Figure S2.2), and in populations with different memory capacity (Figure S3.1b). However, when groups explore the environment more frequently (i.e. when groups make explorative decision more frequently, Figure S4.2, Figure S4.4) and in the resource poor environments where the number of patches is smaller than the number of groups in the environment (Figure S1.2), the presence of a peak largely disappears.

The effect of longer memory (i.e. larger *history* values), exploratory decision, and the number of patches for group to choose from on resource partitioning (Figure 4d-f) closely matches the patterns for patch preferences (Figure 4a-c). One exception is that there is evidence that resource partitioning can peak at intermediate values of history (i.e. resource partitioning in large groups is greatest when groups use more recent memory, Figure 4d). When groups are able to choose among more patches (i.e. when N’_P_ is larger), smaller groups become disproportionately less effective to partition resources than larger groups are (Figure 4f).

### Groups express consistent levels of conflict in preferences among group members, across (larger) group sizes

Overall, groups under the same environmental conditions have a similar percentages of group members whose patch choices match with the most preferred patch in their group (Figure 6). On the other words, living in a group creates a reasonably constant source of conflict in patch choices among individuals, above a given group size. The main driver of conflict is having a larger number of available patches over which groups could choose (N’_P_; full networks vs. ring and random networks in Figure 6, Figure S5.4b). Across all simulations, there is a slight peak in conflict (dip in agreement; also see Figure S3.3, Figure S4.3, Figure S4.6, Figure S5.3), which corresponds to the group size at which preferences begin to be more strongly expressed (Figure 3) but not with the highest rates of resource partitioning (Figure 4). This result is supported by the simulation with higher exploration rates, showing the slight peak in conflict largely disappears when groups explore the environment more frequently (Figure S4.3, Figure S4.6).

**Figure 6.**
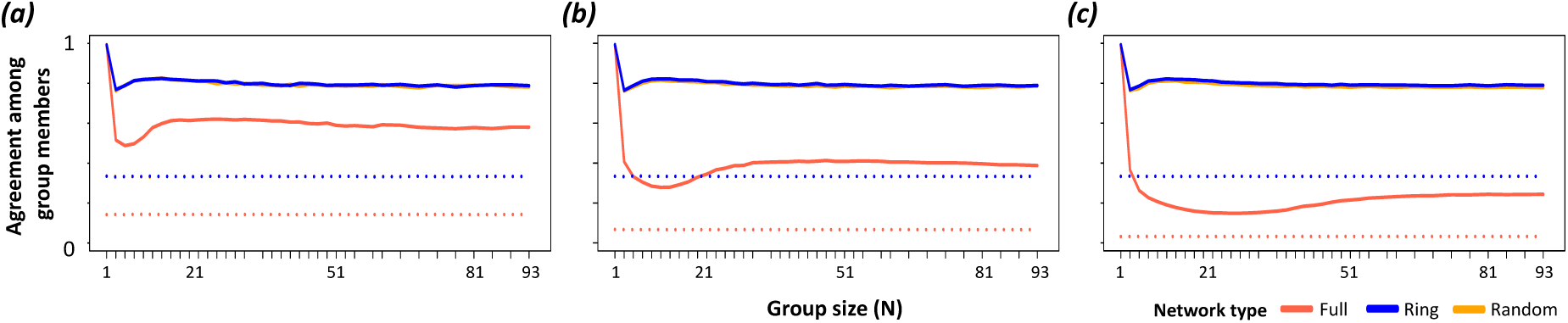
Groups express a relatively constant level of agreement in preferences among group members. Plots show the proportion of group members that share the most common preferences in their group (i.e. agreement), over the last 30 timesteps in simulations with (a) 7, (b) 15, and (c) 31 groups per environment (N_G_). Small groups typically have lower conflict (solitary individuals cannot have any conflict) than larger groups, but conflict remains stable across a large range of group sizes. Across all group sizes, the ability to base decisions on previous experience (solid lines) results in a higher agreement (lower conflict) among group members than when group members make uninformed choices (dashed lines). Groups experience greater conflict when individuals can choose from a larger number of patches (full network) than when they are limited to a subset of patches. The number of groups in the environment (N_G_) does not inherently affect the level of conflicts within groups. Here, 1% of group decisions are explorative (*exploration*=0.01) and individuals remember their own foraging outcome up to three prior timesteps (*history*=3). Colours show full (N’_P_=N_P_), ring (N’_P_=3), and random (N’_P_=3) networks (note that ring and random networks largely overlap).

## Discussion

Our results show that combining short-term memory (only three previous timesteps) with collective decisions-making allows groups of animals to develop strong preferences for where to forage that are distinct from the preferences expressed by other groups. As a consequence of this emergent resource partitioning, individuals in populations of groups can be consistently more successful in accessing resources than individuals in solitary populations. Collective intelligence—in this case negative frequency-dependent learning and information pooling by collectives—allows larger groups to establish stronger preferences than smaller groups and, correspondingly, gain greater fitness benefits. This is especially true when information is more difficult to acquire (i.e. there are more options for each group to choose from). Further, groups can express preferences, and resource partitioning can emerge, even when there is not clear consensus among their members (agreement can be < 0.5 under some environmental conditions). In other words, groups can make adaptive decisions even when group members have diverging preferences. Overall, we propose that this collective sensing of indirect competition can be an additional benefit of group-living, since groups can more accurately estimate which resources are least affected by others, resulting in a more even distribution of groups across the available resources than expected by chance. These results are also robust to modelling decisions, as the patterns are consistent across different environments (Supplemental Information Sections 1 & 2), memory capacity (Supplemental Information Section 3), exploration rates (Supplemental Information Section 4), and the numbers of accessible patches by each groups (Supplemental Information Section 5). Further, once groups in a population establish different preferences, resource partitioning remains stable over time (Supplemental Information Section 6).

### Insights on memory and environmental conditions

Our results reveal that group can establish preferences even when using short-term memory. One perhaps surprising finding is that while using longer-term memory (*history*=7) led to the expression of stronger preferences by groups, these groups could have lower emergent resource partitioning (and lower fitness) than populations of groups using shorter-term (*history*=3) memory. This result highlights an important aspect of considering the broader social (or competitive) environment in individual decision-making. Specifically, the (changing) decisions by other individuals in the population will result in much faster dynamics than the rate at which the environment itself changes. This is also evident from our simulations that start from the highest resource partitioning we detected (Supplemental Information section 6), which show that preferences need to be dynamic to maintain high levels of resource partitioning. As a result, while using a longer history of experiences can facilitate the maintenance of preferences, these preferences are likely to be shaped by increasingly outdated information because the environment (i.e. the decisions of others) changes continuously. Thus, one advantage of collectives is likely that they can aggregate much smaller amounts of (more recent) information into effective decision-making—which is also evidenced by the fact that the peak in resource partitioning is achieved with shorter memory when individuals form larger groups (Figure 4a).

A notable observation is that in a number of environments, the highest resource partitioning was reached by large groups in full networks (e.g. Figures 5 & S3.2). This may be because these environments have the maximum availability of information and large groups can make best use of this information. Reducing the resource availability in the environment also demonstrates the importance of information. In environments with fewer resources than population size (Supplemental Information sections 1 & 2), groups can still express strong preferences but differentiating their preferences from those of other groups can be more challenging (especially when there are more groups than patches). In these environments, solitary individuals and smaller groups perform disproportionately worse than larger groups, likely because they have less information on which to base their decisions. Together, these results highlight how the rate of information gain together with its recency combine to play an important role in shaping the emergence of resource partitioning in animal populations.

### Insights on group living and optimal group sizes

Our study gives insight on several aspects of group living. The first is the contribution of collective sensing of the environment to optimal group sizes [59]. Our model did not implement any costs of making ‘bad’ decisions or of maintaining larger group sizes (e.g. acquiring lower rewards if groups take longer to resolve conflicts in preferences among members [60]). However, we did find that as group size becomes larger, the marginal benefits of increasing group size decreases, and sometimes fitness decreases strongly with increasing group sizes (e.g. when using longer histories, see Figure S3.2). The peaks in resource partitioning support recent predictions that small-to-intermediate group sizes may be able to maximise accuracy of decision-making [61], although this may also depend on the environment as the peak occurred at larger group sizes when groups had to choose amongst more patches (i.e. full networks). If the costs of living in groups also increases with respect to group size (*sensu* [60, 62]), then we expect a strengthening of optimal group size effects.

A further insight is that resource partitioning among groups can emerge even when there are high levels of disagreement within groups. While the benefits in terms of sensing indirect competition increased rapidly before flattening out, consensus costs (i.e. the amount of disagreement among group members, which can be costly [48]) are largely constant across group sizes. One intriguing pattern is that the point where groups begin to express stronger preferences (and individuals begin to gain rewards more often, Figure 5) corresponds to a slight minimum in the level of agreement among group members (Figure 6). This point also corresponds to an increase in the success rate for individuals accessing resources. However, maximum fitness (i.e. highest resource partitioning and reward rates) occur just beyond the peak in disagreement. Together, these results suggest that disagreement may be higher at group sizes that are just below the optimal group size, which could potentially make smaller group sizes unstable. This corresponds with the prediction that stable group sizes should be biased towards larger sizes than the optimal group size [63].

The model also shows that when groups face more choices in their environment (i.e. there are more patches accessible to each group, as represented by full networks versus the ring and random networks), they require a larger size to obtain the same benefits as those living in ‘simpler’ environments (i.e. where there are fewer options to choose from). However, ring and random networks show almost identical patterns, suggesting that differences in how patches are configured in space (or which sets of groups can access them) does not play a critical role in determining the emergence of resource partitioning among groups. Resource richness, by contrast, does not have major effects on these results (Supplemental Information sections 1 & 2). In general, the benefits of being in a large group are slightly lower in resource poor environments than those in resource rich environments, because populations of solitary individuals can more easily partition resources when patches are poorer and because populations across all group sizes find it more difficult to partition resources when there are fewer patches in the environment. Interestingly, in environments where each patch contains smaller amount of resources than group sizes, solitary individuals are better at developing parch preferences in full networks than in other network types, and this effect is stronger in environments with many patches (Figure S2.1). Together, these results suggest that more complex environments (containing more available options or the larger number of groups to indirectly compete with) might select for larger groups, although this effect is slightly weaker in resource-poor environments.

The potential role of collective sensing in shaping group sizes has implications for animals living in harsh environments, where resource availability often fluctuates intensely. For example, during winter or droughts, resources are scarcer and difficult to find, requiring animals to navigate longer distances. Groups of vulturine guineafowl (*Acryllium vulturinum*), a large terrestrial bird endemic to east Africa, substantially increase their home range during periods of drier conditions [64]. While membership to the core group is highly stable, with an optimal group size estimated at 33-37 under intermediate (green, but not wet) conditions [65], these stable groups of vulturine guineafowls have been observed to merge together to form supergroups (group sizes: 65-90) during dry seasons and even larger groups during droughts [64, 66]. Forming such large groups presumably generates substantial benefits through collective intelligence as they navigate unfamiliar environments. Similarly in the superb fairywren (*Malurus cyaneus*), another bird species that forms stable groups, studies conducted in the winter have revealed that they too consistently aggregate into larger groups, which may be linked to navigating larger home ranges [67]. Our models show that larger groups can achieve more consistent access to resources than individuals or smaller groups.

This includes when the environmental signal is more difficult to detect, such as when there are more options to choose from (i.e. N’_P_ is larger), which could occur if or when groups have a larger home range. Further, when groups make explorative decisions more frequently (i.e. higher exploration rates), the optimal group size should also become larger. This is because larger group sizes allow groups to pool foraging outcomes more effectively from these exploratory movements, and living in a larger group and having higher exploration rates might be beneficial when the environment is complex. Multilevel societies—where groups aggregate with other groups, such as in the vulturine guineafowl and super fairy-wrens— could therefore be one way that groups can increase the benefits of collective sensing during periods where conditions are harsh [67], when inaccurate decision-making might cost disproportionately more time and energy (e.g. opportunity costs), and in situations where individuals live in fluctuating environments.

### Assumptions of the model

Our model has some key assumptions. The main assumption is that individuals can only acquire information about indirect competition. In reality, individuals will often have access to many other cues, including through direct interactions with other groups at patches. Our model can also be informative about such situations by reframing the information memorised by individuals as the patch being a good patch vs. a bad patch (*sensu* [68]). Determinants of patch quality could include both direct (e.g. competitive interactions) and indirect cues (whether they foraged successfully or not). The model could also be extended to allow individuals to acquire a continuum of information between these two binary states—such as differentiating between acquiring food versus not acquiring food while also experiencing competitive interactions. However, such an implementation is unlikely to change the core results of our study—that by pooling information, populations of groups can better partition resources.

A second assumption is that individuals only show preferences for patches where they foraged successfully, and not avoidance for patches where they were not successful. Thus, previously not visited patches and unsuccessful patches are considered equal. This has some degree of realism, for example a group that moves through a depleted patch may not be able to perceive that this patch previously contained food. However, if they could actively detect depletion, and memorise this information, then this information could be encoded in a way that reduces the perceived quality of each patch. This could have some adaptive value, as it would decrease the chance that a group visits a patch for which it has information about previous depletion and, correspondingly, increase the chances of visiting completely unexplored patches (e.g. when making exploratory decisions). We thus predict that groups should become better at developing preferences, and populations should become better at resource partitioning, if individuals can sense patch depletion.

A third assumption is that food resources at a patch are distributed equally across groups visiting the patch at the same time. One alternative would be to allocate the food at random. Doing so should simply act to increase the differences in the perceived quality of patches among the visiting groups, and thus increase the rate at which preferences (and subsequently resource partitioning) arise. A second alternative is to define one group to receive all of the food (either randomly or following a dominance hierarchy among groups). Doing so, however, would revert the model towards the same condition as single individuals, as all group members would have exactly the same information. Whether groups can still perform better than individuals might then depend on *how* consensus is reached among group members. Several forms of collective decision-making exist (e.g. a single leader, a sub-majority, or a quorum; [26, 48]), and future work may use our model to explore the consequences that these have on the emergent patterns at the population level.

Finally, a fourth assumption of our model is that all group members are ‘equal’, and thus that they are equally likely to gain a food resource when multiple groups visit a patch. In reality, access to resources is often stratified among individuals according to dominance hierarchies, such that a subset of individuals more frequently acquires food and another subset more rarely acquires food. Previous work has demonstrated that these dynamics can affect the timing of departure of groups from patches [69], and that dominant individuals who have more potential to gain from rich patches can lead their group to those patches [70]. However, how competition within groups shapes the ability for collective sensing of indirect competition, and how this affects the distribution of groups over space when there are many patches, remains unknown.

### Insights on territoriality and specialisation across species

Our study shows that memory—in our case very short-term or limited memory—can enable populations of group-living animals to more effectively establish distinct spatial ranges (here by specialising on one food patch) than populations of solitary animals. These results extend previous theoretical and empirical studies that found memory can play a key role in individuals partitioning space [46] or resources [32] through negative frequency-dependent learning. Specifically, collective sensing of resource gradients created by indirect competition could potentially play a role in driving the evolution of territoriality. Our study captures the first potential steps in this process. Once individuals (or groups) begin gaining more benefits by developing preferred patches relative to the background expectation of the environment, this should increase their perceived value of the resource, thereby increasing their incentives to begin defending them. Supplementary Information section 6 demonstrates that the dynamics introduced by the social environment requires groups to continuously update their preferences, thereby introducing a cost that could be offset by resource defense. Defending a preferred resource patch could also drive other contingent benefits. For example, limiting the number of patches a group accesses can increase their familiarity with the environment, thereby allowing groups to exploit patches more effectively while have fewer conflicts in preferences among members [71–73]. For example, familiarity can reduce navigational errors and reducing decision-making time, thereby reducing costs in terms of time investment [48], and, in turn, making them more efficient at exploiting resources from their environment (producing a positive feedback). Thus, the benefits derived from resource partitioning can promote active defense of the preferred resources from other competitors, even when similar resources might be available elsewhere in the environment. Thus, we propose that one pathway through which territoriality could emerge is via a process of self-organisation— initially facilitated by negative frequency-dependent learning of indirect competition for resources—that is enhanced in group-living animals.

While our model is focused on populations consisting of competing groups from one population of one species, our results may further be extended to consider the role of collective behaviour in the evolution of specialisations across species. Feedbacks or social interactions among sets of individuals could stimulate opportunities for them to collectively develop preferences for certain type of resources or foraging strategies. For example, Aplin et al. [40] showed that great tit (*Parus major*) groups developed the preference for experimentally-introduced solutions over the alternative solutions with equal rewards, when individuals learnt the experimentally-introduced foraging cues through social learning.

Extending this finding across species, a study cross-fostering great tits and blue tits nestlings showed that individuals’ foraging niche can be shaped by social information from their rearing parents [74, 75]. As social learning allows information to transmit faster within and across generations, emergent preferences arising from negative frequency-dependent learning within sets of individuals in a population (e.g. different species) could rapidly become entrenched over generations, even before any morphological differences are present. Thus, various forms of specialisation (from territoriality to the evolution of ecological niches) could potentially emerge even in the absence of strong selection on intergroup resource defence or on pre-existing variation in morphological traits, through a process of self-organisation at the population-level that is driven by negative frequency-dependent learning and collective intelligence.

## Acknowledgments

We thank Mauricio Cantor, Eli Strauss, and the Farine lab for feedbacks and comments.

## Author contributions

DRF coded the core model and MO extended the model. MO wrote the first draft, and both authors revised the manuscript.

