## Supplemental Information for "Collective intelligence facilitates emergent resource partitioning through frequency dependent learning"

### Section 1. The effect of having fewer patches in the environment

We tested whether the results we observed would be consistent in resource-poor environments (i.e. where competition for resources is greater), when modifying the number of patches in the environment. We set the number of patches in the environment to be a maximum of two thirds of the number of groups (i.e. $N_P=\lfloor 2N_G/3 \rfloor$). For example, if there are 7 groups then we created four patches (2*7/3=4.6, which rounds down to 4). Each patch still has the same amount of resources as group size (N), per the main simulations, but because there are fewer patches, the total amount of food in the environment is lower.

Overall, the results were consistent with the environmental condition simulated in the main text (i.e. where $N_P=N_G$; Figure 3,4,5,6), despite the foraging success being lower overall (comparing to the condition with $N_P=N_G$; Figure 5). Specifically, groups develop stronger patch preferences (Figure S1.1), with individuals gaining greater foraging success when foraging as groups (Figure S1.2) and with groups experiencing consistent level of disagreement among group members (Figure S1.3).

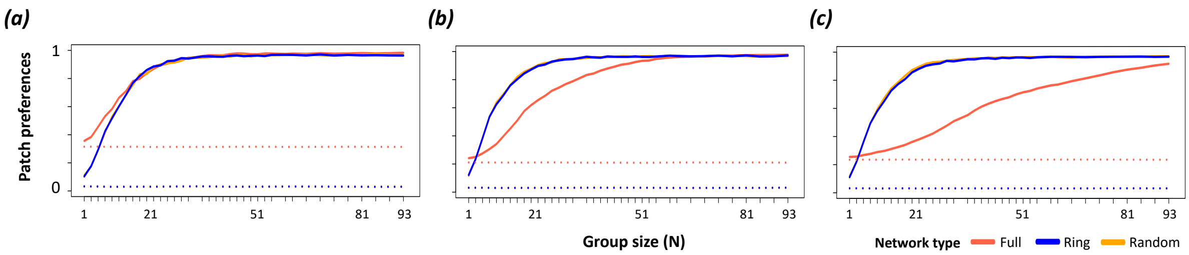


**Figure S1.1. Groups can develop strong patch preferences.** These plots show the patch preferences (1-normalised entropy in patch use within groups) over the last 30 timesteps in simulations with (a) 7, (b) 15, and (c) 31 groups per environment ($N_G$) where the number of patches in the environment ($N_P$) is smaller than the number of groups in the environment ($N_P<N_G$). Patch preference is higher in groups than solitary individuals and increases with group size (x axis) when individuals use prior experience (*history*=3, solid lines) to choose foraging patches (relative to making explorative patch choices at all timesteps, dotted lines, *history*=0). While only large groups develop preferences when they can access many patches [e.g. $N_P$=31 in the full network, orange line, in (c)], even small groups (e.g. N=11) can develop strong preferences when they are restricted in the number of patches that they can access (i.e. ring and random networks, where $N'_P<N_G$). Note that the maximum patch preferences are generally slightly less than 1 because simulations were programmed such that groups make explorative decisions in 1% of timesteps (*exploration*=0.01). Colours show full ($N'_P=N_P$), ring ($N'_P$=3), and random ($N'_P$=3) networks (note that ring and random networks largely overlap and may not both be visible).

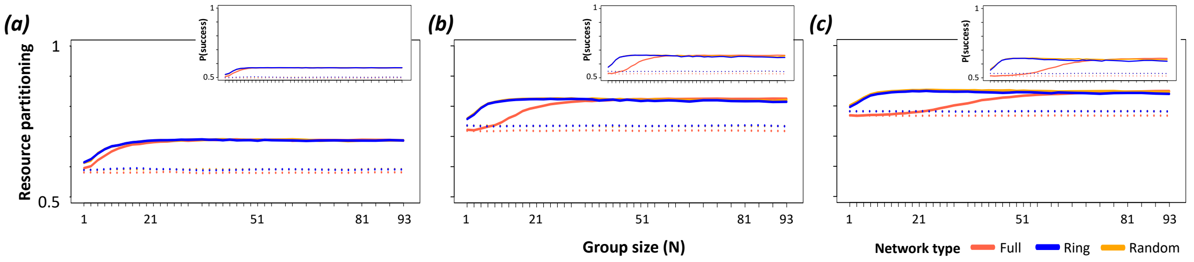


**Figure S1.2. Groups show stronger and consistent resource partitioning across the population, consistently individuals in groups with stronger preferences experience higher foraging rates.** These plots show the resource partitioning (normalised entropy in patch use across groups; main figures) and the frequency of success in foraging (insets of panels) over the last 30 timesteps in simulations with (a) 7, (b) 15, and (c) 31 groups per environment, $N_G$. Stronger preferences (see Figure 3) lead to higher individual foraging rates, and therefore greater partitioning of resources (resource use specialisation) among groups than among solitary individuals. Colours show full (**$N'_P=N_P$**), ring ($N'_P=3$), and random ($N'_P=3$) networks (note that ring and random networks largely overlap).

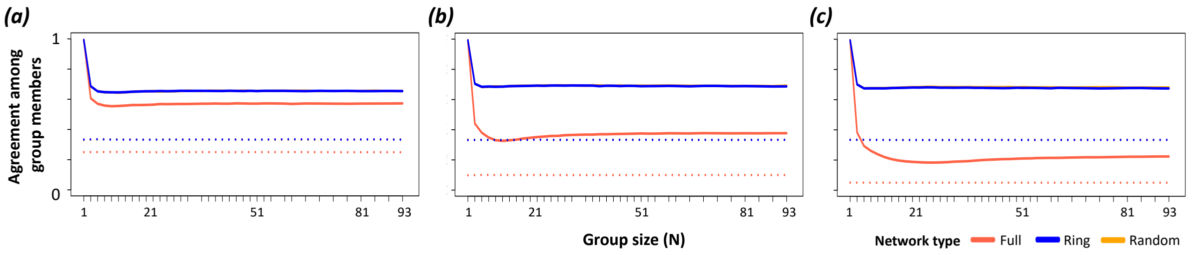


**Figure S1.3. Groups express a relatively constant level of agreement in preferences among group members.** Small groups typically have lower conflict (solitary individuals cannot have any conflict) than larger groups, but conflict remains stable across a large range of group sizes. Across all group sizes, the ability to base decisions on previous experience (solid lines, *history*=3) results in a higher agreement (lower conflict) among group members than when group members choose patches exploratively (dashed lines, *history*=0). Groups experience greater conflict when individuals can choose from a larger number of patches (full network) than when they are limited to a subset of patches. The number of groups in the environment ($N_G$) does not inherently affect the level of conflicts within groups. Here, 1% of collective decisions were explorative (*exploration*=0.01) and individuals had access to their own foraging outcome up to three prior timesteps (*history*=3). Colours show full ($N'_P=N_P$), ring ($N'_P=3$), and random ($N'_P=3$) networks (note that ring and random networks largely overlap).

### Section 2. The effect of insufficient number of food resources per patch

We tested whether the results we observed would be consistent in resource-poor environments (i.e. where competition for resources is greater), when modifying the amount of food resources per patch. We set each simulated patch in the environment to contain approximately two thirds of group sizes (i.e. patch size is $\lfloor N*2/3 \rfloor$, where N is group size). For example, if there are 7 groups of 15 individuals, the simulated environment contains 7 patches where each patch has 10 resources (15*2/3=10). The number of patches in the environment stayed the same as the number of simulated groups (i.e. $N_P=N_G$), per the main simulations, but because there are fewer resources per patch, the total amount of food in the environment is lower.

Overall, the results were consistent with the environmental condition simulated in the main text (i.e. where patch sizes were equal to group sizes; Figure 3,4,5,6), despite the foraging success being lower overall (comparing to the condition where patch sizes were equal to group sizes; Figure 5). Specifically, groups develop stronger patch preferences (Figure S2.1), with individuals gaining greater foraging success when foraging as groups (Figure S2.2) and with groups experiencing consistent level of disagreement among group members (Figure S2.3). Note that, since the patch size is $\lfloor N*2/3 \rfloor$ (i.e. smaller than group sizes), solitary individuals were always unsuccessful as default, but as group size increases (e.g. group size of 3) individuals became more successful.

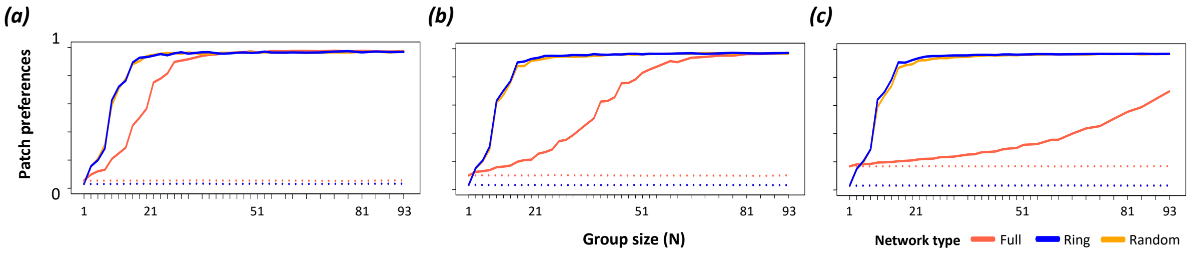


**Figure S2.1. Groups can develop strong patch preferences.** These plots show the patch preferences (1-normalised entropy in patch use within groups) over the last 30 timesteps in simulations with (a) 7, (b) 15, and (c) 31 groups per environment ($N_G$) where patch sizes are smaller than group sizes (patch size<N). Patch preference is higher in groups than solitary individuals and increases with group size (x axis) when individuals use prior experience (*history*=3, solid lines) to choose foraging patches (relative to making explorative patch choices at all timesteps, dotted lines, *history*=0). While only large groups develop preferences when they can access many patches [e.g. $N_P$ = 31 in the full network, orange line, in (c)], even small groups (e.g. N=11) can develop strong preferences when they are restricted in the number of patches that they can access (i.e. ring and random networks, where $N'_P<N_G$). Note that the maximum patch preferences is generally slightly less than 1 because simulations were programmed such that groups make explorative decisions in 1% of timesteps (*exploration*=0.01). Colours show full ($N'_P=N_P$), ring ($N'_P=3$), and random ($N'_P=3$) networks (note that ring and random networks largely overlap and may not both be visible).

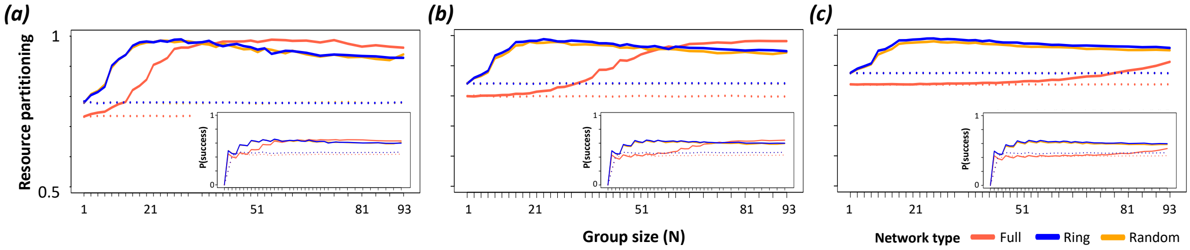


**Figure S2.2. Groups show stronger and consistent resource partitioning across the population, consistently individuals in groups with stronger preferences experience higher foraging rates.** These plots show the resource partitioning (normalised entropy in patch use across groups; main figures) and the frequency of success in foraging (insets of panels) over the last 30 timesteps in simulations with (a) 7, (b) 15, and (c) 31 groups per environment. Stronger preferences (see Figure 3) lead to higher individual rewards, and therefore greater partitioning of resources (resource use specialisation) among groups than among solitary individuals. Colours show full ($N'_P=N_P$), ring ($N'_P=3$), and random ($N'_P=3$) networks (note that ring and random networks largely overlap).

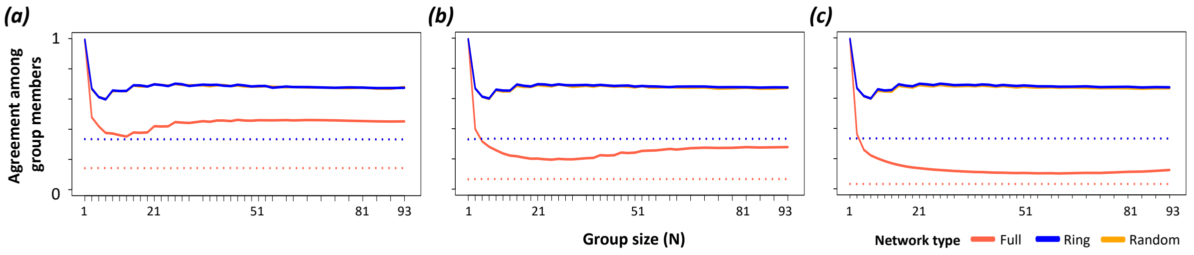


**Figure S2.3. Groups express a relatively constant level of agreement in preferences among group members.** Small groups typically have lower conflict (solitary individuals cannot have any conflict) than larger groups, but conflict remains stable across a large range of group sizes. Across all group sizes, the ability to base decisions on previous experience (solid lines, *history*=3) results in a higher agreement (lower conflict) among group members than when group members choose patches exploratively (dashed lines, *history*=0). Groups experience greater conflict when individuals can choose from a larger number of patches (full network) than when they are limited to a subset of patches. The number of groups in the environment ($N_G$) does not inherently affect the level of conflicts within groups. Here, 1% of collective decisions were explorative (*exploration*=0.01) and individuals had access to their own foraging outcome up to three prior timesteps (*history*=3). Colours show full ($N'_P$=$N_P$), ring ($N'_P$=3), and random ($N'_P$=3) networks (note that ring and random networks largely overlap).

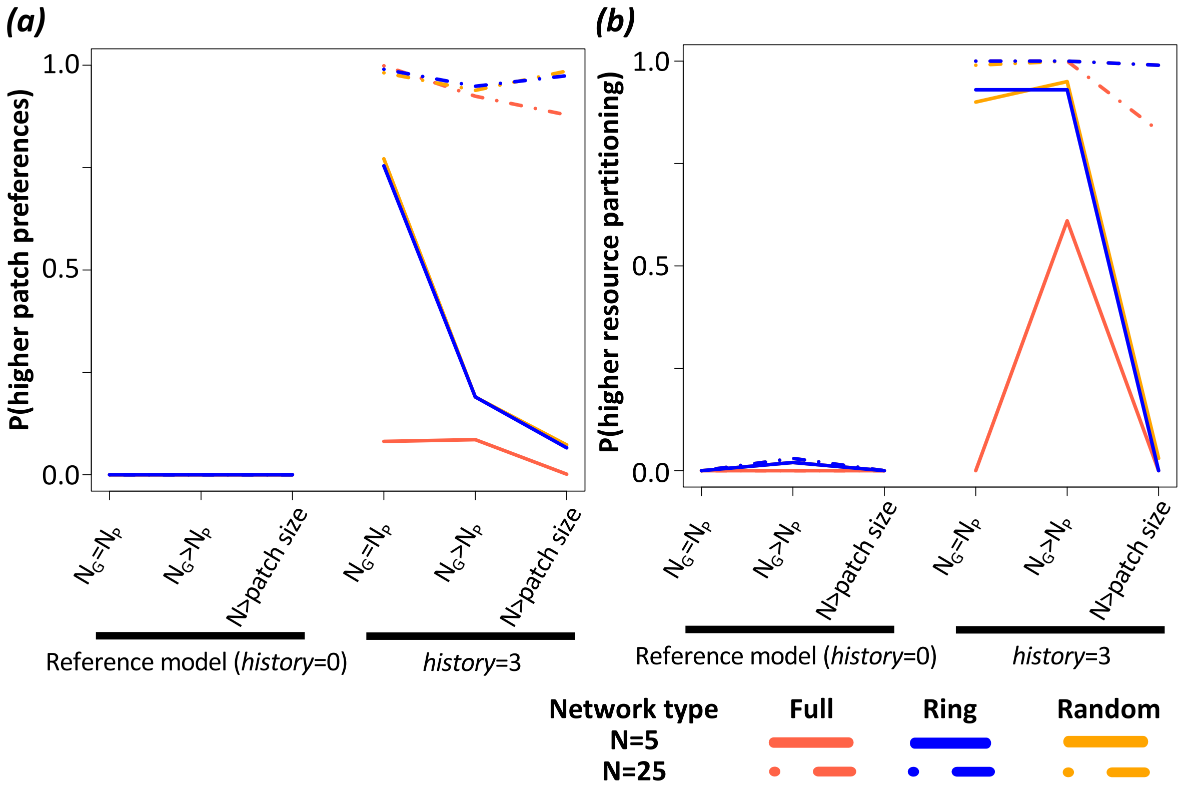


**Figure S2.4. The total amount of resources due to fewer number of patches and reduced number of resources per patch affects the establishment of preferences, emergence of resource partitioning, consequently individuals foraging success, and the level of agreement among group members.** Plots show the effects of resource amounts and distributions on four metrics: patch preferences (*a*) and resource partitioning among groups (*b*), individuals foraging rates (c), the agreement among group members (d). The y axis shows the tendency of the simulation with the given parameter set to express a higher metric value that is higher than or equal to the median of the default parameter set (see methods). Colours show full ($N'_P=N_P$), ring, and random networks (note that results for the different networks often overlap with each other).

### Section 3. The effect of increasing memory

We tested the sensitivity of the memory capacity by modifying *history* parameter. When memory capacity is larger, groups develop stronger patch preferences. Nevertheless, the overall results remain qualitatively similar. Specifically, groups develop stronger patch preferences and individuals in cohesive groups successfully acquire resource more often than solitary individuals, and living in a group creates a reasonably constant source of conflict in patch choices among individuals, above a given group size.

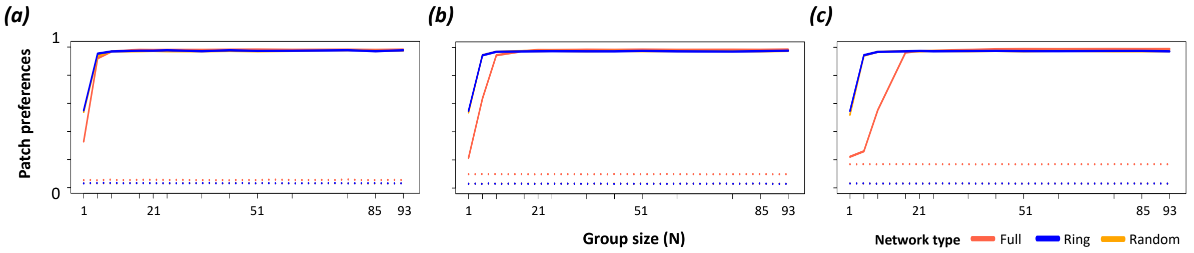


**Figure S3.1. Groups can develop strong patch preferences.** These plots show the patch preferences (1-normalised entropy in patch use within groups) over the last 30 timesteps in simulations with (a) 7, (b) 15, and (c) 31 groups per environment ($N_G$). Patch preference is higher in groups than solitary individuals and increases with group size (x axis) when individuals use prior experience (***history*=7**, solid lines) to choose foraging patches (relative to making uninformed patch choices in all timesteps, dotted lines, *history*=0). While only large groups develop preferences when they can access many patches [e.g. $N_P$=31 in the full network, orange line, in (c)], even small groups (e.g. N=11) can develop strong preferences when they are restricted in the number of patches that they can access (i.e. ring and random networks, where $N'_P < N_G$). Note that the maximum patch preferences is generally slightly less than 1 because simulations were programmed such that groups make explorative decisions in 1% of timesteps (*exploration*=0.01). Colours show full ($N'_P$=$N_P$), ring ($N'_P$=3), and random ($N'_P$=3) networks (note that ring and random networks largely overlap and may not both be visible).

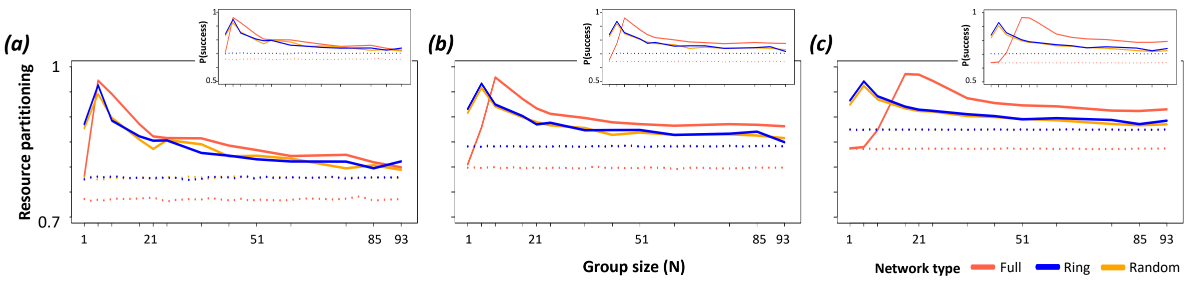


**Figure S3.2. Groups show stronger and consistent resource partitioning across the population, consistently individuals in groups with stronger preferences experience higher foraging rates.** These plots show the resource partitioning (normalised entropy in patch use across groups; main figures) and the frequency of success in foraging (insets) over the last 30 timesteps in simulations with (a) 7, (b) 15, and (c) 31 groups per environment, $N_G$. Stronger preferences (see Figure 3) lead to higher individual foraging rates, and therefore greater partitioning of resources (resource use specialisation) among groups than among solitary individuals. Colours show full ($N'_P$=$N_P$), ring ($N'_P$=3), and random ($N'_P$=3) networks (note that ring and random networks largely overlap).

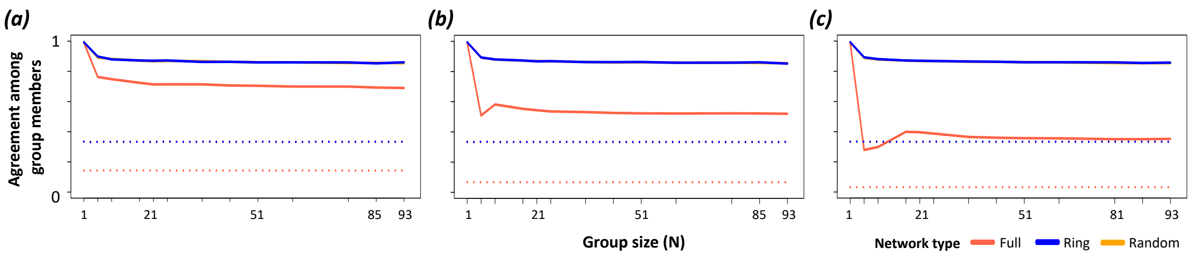


**Figure S3.3. Groups express a relatively constant level of agreement in preferences among group members.** Small groups typically have lower conflict (solitary individuals cannot have any conflict) than larger groups, but conflict remains stable across a large range of group sizes. Across all group sizes, the ability to base decisions on previous experience (solid lines, ***history***=7) results in a higher agreement (lower conflict) among group members than when group members choose patches exploratively (dashed lines, *history*=0). Groups experience greater conflict when individuals can choose from a larger number of patches (full network) than when they are limited to a subset of patches. The number of groups in the environment ($N_G$) does not inherently affect the level of conflicts within groups. Here, 1% of collective decisions were explorative (*exploration*=0.01) and individuals had access to their own foraging outcome up to three prior timesteps (*history*=7). Colours show full ($N'_P$=$N_P$), ring ($N'_P$=3), and random ($N'_P$=3) networks (note that ring and random networks largely overlap).

### Section 4. The effect of exploration rate

We tested the sensitivity of exploration rates. When groups make explorative decisions more frequently (i.e. higher exploration rates), the optimal group size should also become larger. Nevertheless, the overall results remain the similar, that is groups develop stronger patch preferences and individuals in cohesive groups successfully acquire resource more often than solitary individuals, and living in a group creates a reasonably constant source of conflict in patch choices among individuals, above a given group size.

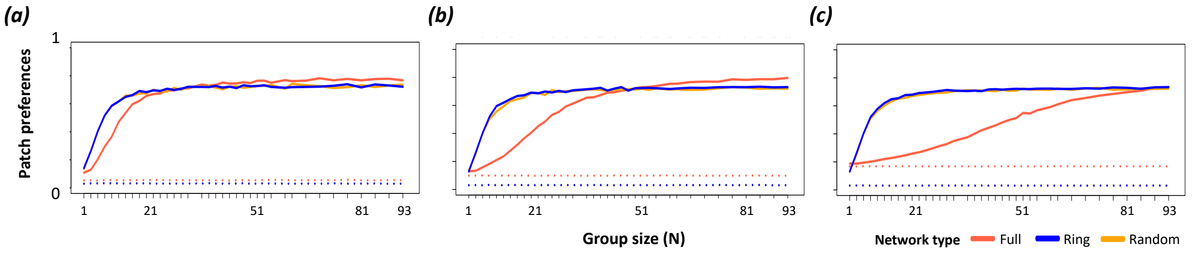


**Figure S4.1. Groups can develop strong patch preferences.** These plots show the patch preferences (1-normalised entropy in patch use within groups) over the last 30 timesteps in simulations with (a) 7, (b) 15, and (c) 31 groups per environment ($N_G$). Patch preference is higher in groups than solitary individuals and increases with group size (x axis) when individuals use prior experience (*history*=3, solid lines) to choose foraging patches (relative to making explorative patch choices in all timesteps, dotted lines, *history*=0). While only large groups develop preferences when they can access many patches [(e.g. $N_P$=31 in the full network, orange line, in (c)], even small groups (e.g. N=11) can develop strong preferences when they are restricted in the number of patches that they can access (i.e. ring and random networks, where $N'_P<N_G$). Note that groups make explorative decisions in 10% of timesteps (***exploration*=0.1**). Colours show full ($N'_P=N_P$), ring ($N'_P$=3), and random ($N'_P$=3) networks (note that ring and random networks largely overlap and may not both be visible).

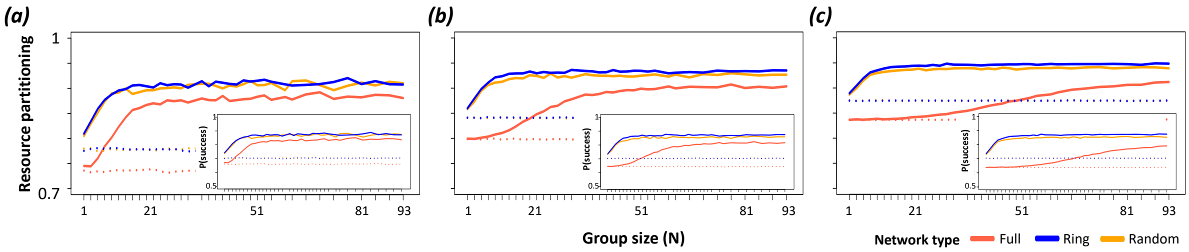


**Figure S4.2. Groups show stronger and consistent resource partitioning across the population, consistently individuals in groups with stronger preferences experience higher foraging rates.** These plots show the resource partitioning (normalised entropy in patch use across groups; main figures) and the frequency of success in foraging over the last 30 timesteps in simulations with (a) 7, (b) 15, and (c) 31 groups per environment, $N_G$. Stronger preferences (see Figure 3) lead to higher individual foraging rates (insets), and therefore greater partitioning of resources (resource use specialisation) among groups than among solitary individuals. Colours show full ($N'_P=N_P$), ring ($N'_P=3$), and random ($N'_P=3$) networks (note that ring and random networks largely overlap).

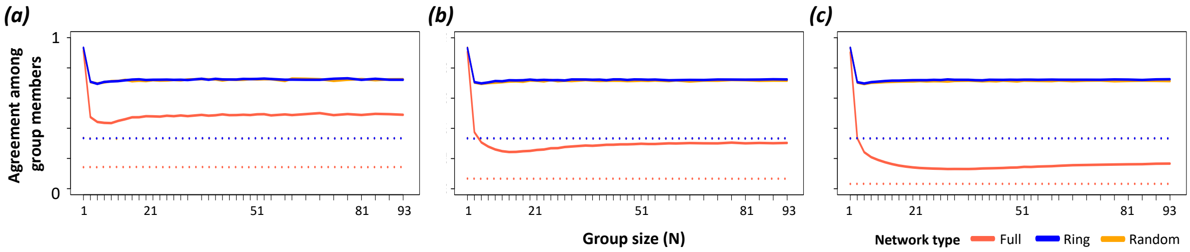


**Figure S4.3. Groups express a relatively constant level of agreement in preferences among group members.** Small groups typically have lower conflict (solitary individuals cannot have any conflict) than larger groups, but conflict remains stable across a large range of group sizes. Across all group sizes, the ability to base decisions on previous experience (solid lines, ***history*=3**) results in a higher agreement (lower conflict) among group members than when group members choose patches exploratively (dashed lines, ***history*=0**). Groups experience greater conflict when individuals can choose from a larger number of patches (full network) than when they are limited to a subset of patches. The number of groups in the environment ($N_G$) does not inherently affect the level of conflicts within groups. Here, 10% of collective decisions were explorative (***exploration*=0.1**) and individuals had access to their own foraging outcome up to three prior timesteps (*history*=3). Colours show full ($N'_P=N_P$), ring ($N'_P=3$), and random ($N'_P=3$) networks (note that ring and random networks largely overlap).

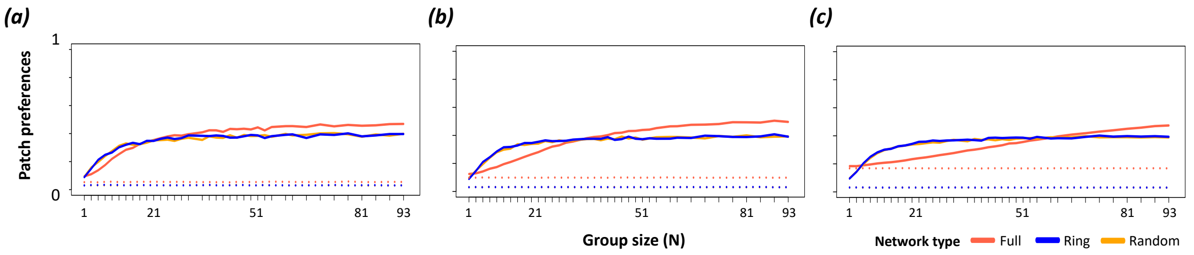


**Figure S4.4. Groups can develop strong patch preferences.** These plots show the patch preferences (1-normalised entropy in patch use within groups) over the last 30 timesteps in simulations with (a) 7, (b) 15, and (c) 31 groups per environment ($N_G$). Patch preference is higher in groups than solitary individuals and increases with group size (x axis) when individuals use prior experience (*history*=3, solid lines) to choose foraging patches (relative to making explorative patch choices in all timesteps, dotted lines, *history*=0). While only large groups develop preferences when they can access many patches [e.g. $N_P$=31 in the full network, orange line, in (c)), even small groups (e.g. N=11) can develop strong preferences when they are restricted in the number of patches that they can access (i.e. ring and random networks, where $N'_P$<$N_G$ as $N_G$ increases). Note that groups make explorative decisions in 25% of timesteps (***exploration*=0.25**). Colours show full ($N'_P$=$N_P$), ring ($N'_P$=3), and random ($N'_P$=3) networks (note that ring and random networks largely overlap and may not both be visible).

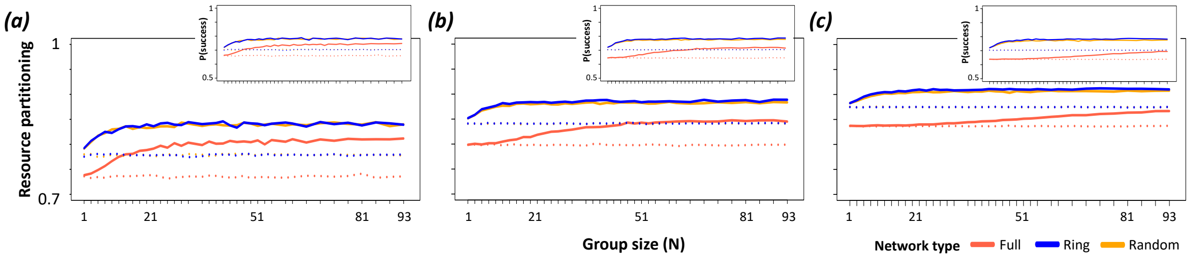


**Figure S4.5. Groups show stronger and consistent resource partitioning across the population, consistently individuals in groups with stronger preferences experience higher foraging rates.** These plots show the resource partitioning (normalised entropy in patch use across groups; main figures) and the frequency of success in foraging over the last 30 timesteps in simulations with (a) 7, (b) 15, and (c) 31 groups per environment, $N_G$. Stronger preferences (see Figure 3) lead to higher individual foraging rates (insets), and therefore greater partitioning of resources (resource use specialisation) among groups than among solitary individuals. Colours show full ($N'_P$=$N_P$), ring ($N'_P$=3), and random ($N'_P$=3) networks (note that ring and random networks largely overlap).

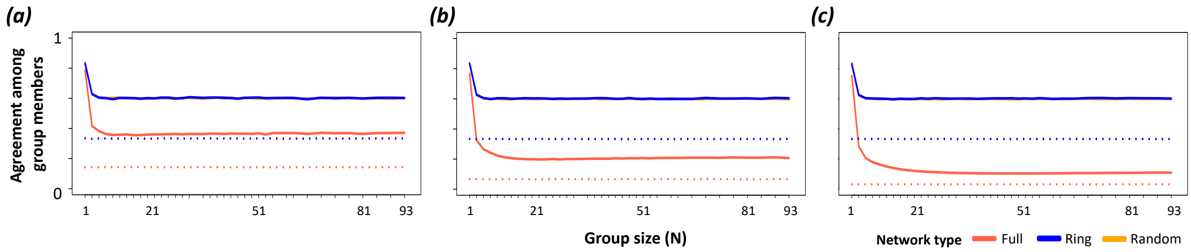


**Figure S4.6. Groups express a relatively constant level of agreement in preferences among group members.** Small groups typically have lower conflict (solitary individuals cannot have any conflict) than larger groups, but conflict remains stable across a large range of group sizes. Across all group sizes, the ability to base decisions on previous experience (solid lines, *history*=3) results in a higher agreement (lower conflict) among group members than when group members choose patches exploratively (dashed lines, *history*=0). Groups experience greater conflict when individuals can choose from a larger number of patches (full network) than when they are limited to a subset of patches. The number of groups in the environment ($N_G$) does not inherently affect the level of conflicts within groups. Here, 25% of collective decisions were explorative (***exploration*=0.25**) and individuals had access to their own foraging outcome up to three prior timesteps (*history*=3). Colours show full ($N'_P=N_P$), ring ($N'_P=3$), and random ($N'_P=3$) networks (note that ring and random networks largely overlap).

#### Section 5. The effect of different number of patches that groups can choose from when groups are spatially restricted

We tested the sensitivity of the different numbers of accessible patches by groups in models where they are spatially restricted (i.e. they can access a subset of all patches). As the number of available patches to sample for each group increases (***degree*=5** vs. ***degree*=3)**, the ability for groups to develop a preference weakens. Nevertheless, the overall results remain the similar, that is groups develop stronger patch preferences and individuals in cohesive groups successfully acquire resource more often than solitary individuals, and living in a group creates a reasonably constant source of conflict in patch choices among individuals, above a given group size.

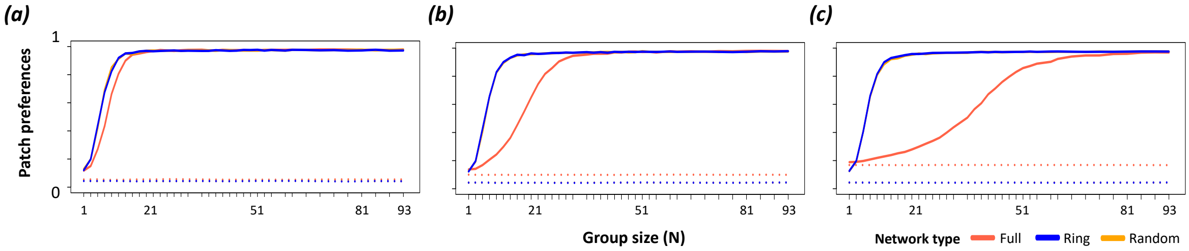


**Figure S5.1. Groups can develop strong patch preferences.** These plots show the patch preferences (1-normalised entropy in patch use within groups) over the last 30 timesteps in simulations with (a) 7, (b) 15, and (c) 31 groups per environment ($N_G$). Patch preference is higher in groups than solitary individuals and increases with group size (x axis) when individuals use prior experience (*history*=3, solid lines) to choose foraging patches (relative to explorative patch choices, dotted lines, *history*=0). While only large groups develop preferences when they can access many patches [e.g. $N_P$=31 in the full network, orange line, in (c)], even small groups (e.g. N=11) can develop strong preferences when they are restricted in the number of patches that they can access (i.e. ring and random networks, where $N'_P<N_G$). Note that the maximum patch preferences is generally slightly less than 1 because simulations were programmed such that groups make explorative decisions in 1% of timesteps (*exploration*=0.01). Colours show full (**$N'_P=N_P$**), ring (**$N'_P$=5**), and random (**$N'_P$=5**) networks (note that ring and random networks largely overlap and may not both be visible).

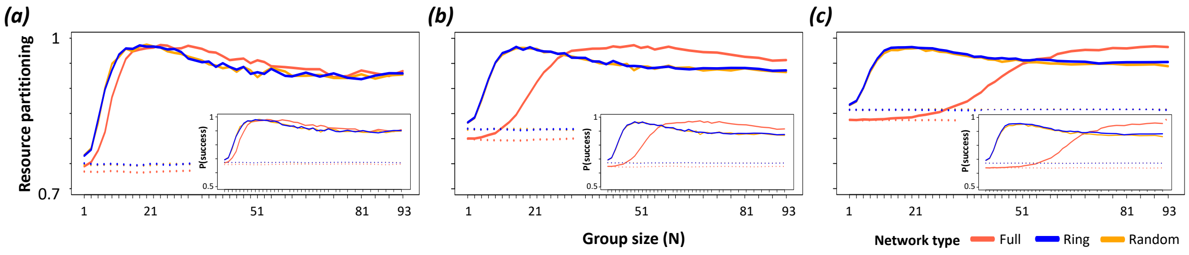


**Figure S5.2. Groups show stronger and consistent resource partitioning across the population, consistently individuals in groups with stronger preferences experience higher foraging rates.** These plots show the resource partitioning (normalised entropy in patch use across groups; main figures) and the frequency of success in foraging over the last 30 timesteps in simulations with (a) 7, (b) 15, and (c) 31 groups per environment, $N_G$. Stronger preferences (see Figure 3) lead to higher individual foraging rates (insets), and therefore greater partitioning of resources (resource use specialisation) among groups than among solitary individuals. Colours show full (**$N'_P=N_P$**), ring (**$N'_P=5$**), and random (**$N'_P=5$**) networks (note that ring and random networks largely overlap).

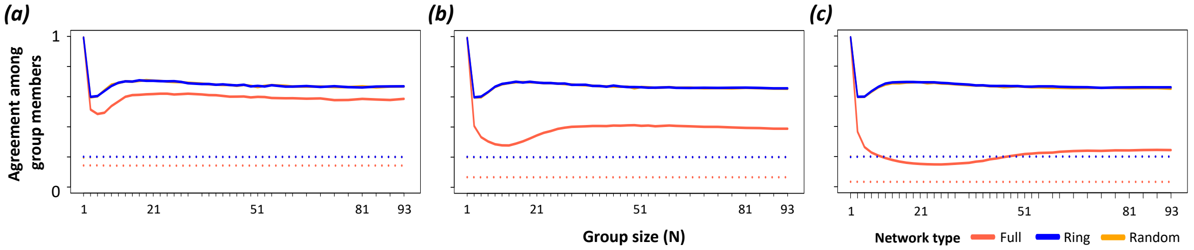


**Figure S5.3. Groups express a relatively constant level of agreement in preferences among group members.** Small groups typically have lower conflict (solitary individuals cannot have any conflict) than larger groups, but conflict remains stable across a large range of group sizes. Across all group sizes, the ability to base decisions on previous experience (solid lines, ***history*=3**) results in a higher agreement (lower conflict) among group members than when group members choose patches exploratively (dashed lines, ***history*=0**). Groups experience greater conflict when individuals can choose from a larger number of patches (full network) than when they are limited to a subset of patches. The number of groups in the environment ($N_G$) does not inherently affect the level of conflicts within groups. Here, 1% of collective decisions were explorative (*exploration*=0.01) and individuals had access to their own foraging outcome up to three prior timesteps (*history*=3). Colours show full (**$N'_P=N_P$**), ring (**$N'_P=5$**), and random (**$N'_P=5$**) networks (note that ring and random networks largely overlap).

### Section 6. Stability of patch preferences across timesteps

We tested whether patch preferences developed by groups remain stable across timesteps. First, we ran the simulation up to 300 timesteps and determined the focal metrics for each iteration for each simulation from the last 30 timesteps (271-300 timesteps), and identified the iteration where groups showed the highest patch preferences. Expanding this focal simulation up to 350 timesteps and repeating this expansion 100 iterations, we determined the four metrics for each iteration over the last 30 timesteps (321-350 timesteps) and quantified how much the four metrics diverge when starting from the same conditions (the identical parameter set and sharing the same foraging history). Overall, groups maintain extremely high patch preferences and resource partitioning (and, consequently, individuals foraging rates remain higher). We note that even though the initial condition is identical across iterations, slight variations in metrics emerge after the additional 50 timesteps. This indicates that emerging resource specialisation and partitioning via collective sensing through negative frequency-dependent learning and information pooling is a dynamic process, and the transitionary nature of the dynamics requires groups to keep updating information sampled from the recent environment.

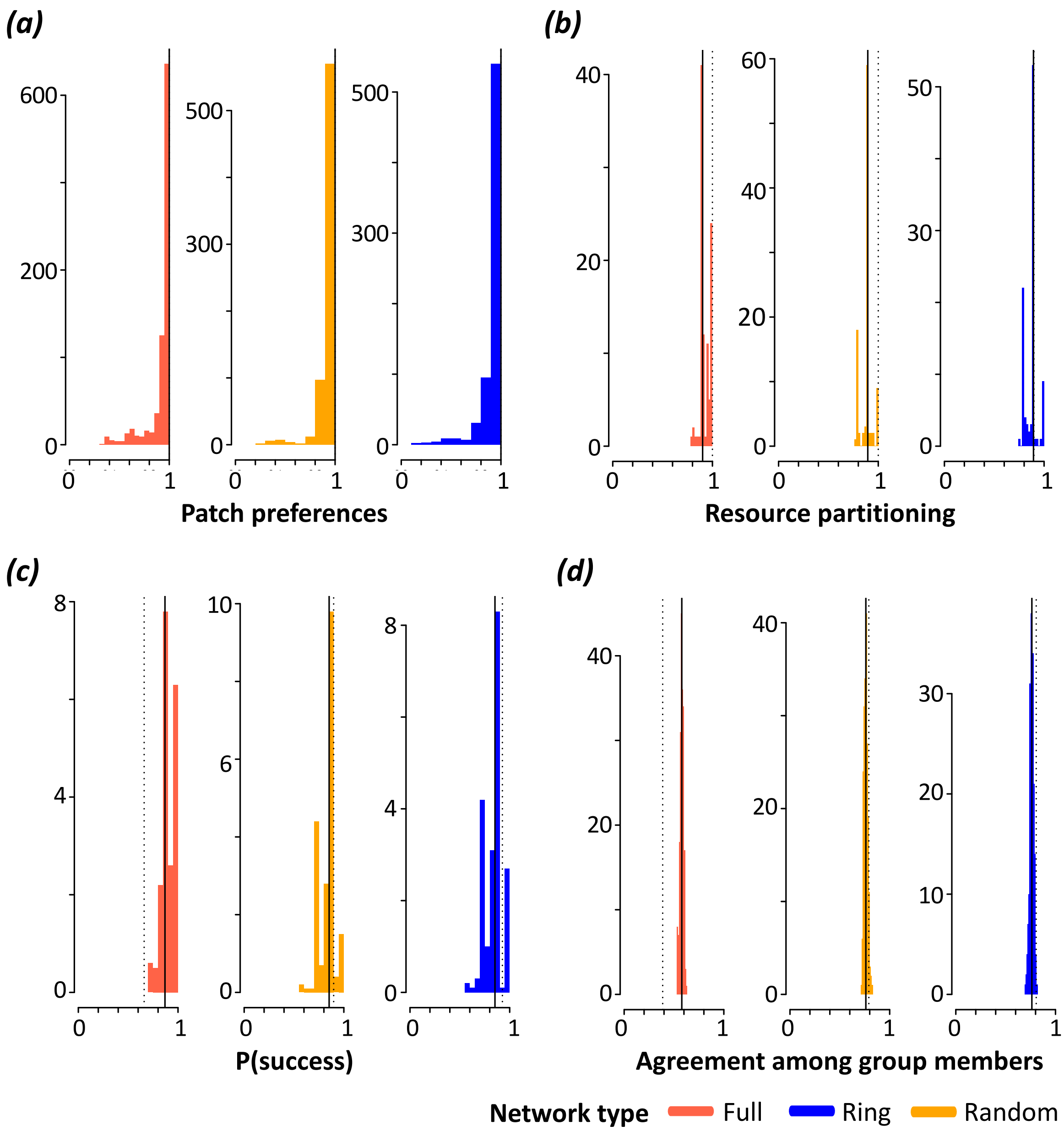


**Figure S6.1. Developed preferences for specific patches by groups can be stable.** The x-axes show the range of the focal metric: patch preferences (a), resource partitioning (b), foraging success at individual level (c), and the agreement among group members (d). The y-axes show the frequency that groups expressed the corresponding metric values, over the last 30 timesteps from the additional timesteps (i.e. 321-350 timesteps) from simulations with 7 groups (**$N_G$=7**) of 25 (**N=25**) and whose 1% of decision-making was explorative (***exploration*=0.01**). Each simulated individual has a memory capacity of 3 (***history*=3**). The solid black line shows the median value of the corresponding metric from 321-350 timesteps, and the dotted black line shows the metric value before simulation was extended (i.e. the corresponding metric from the iteration with the highest normalized patch preferences over 271-300 timesteps). Colours show full ($N_P$=$N_G$), ring (**$N'_P$=3**), and random (**$N'_P$=3**) networks (note that ring and random networks largely overlap).
